# Sex differences in alternative reproductive tactics in response to predation risk in tree crickets

**DOI:** 10.1101/848630

**Authors:** Viraj R. Torsekar, Rohini Balakrishnan

## Abstract

Predation risk has been hypothesised to drive the evolution of alternative mate-search strategies, but few empirical studies have examined this. In crickets, mate-search involves acoustic signalling by males and acoustic-mediated movement by females. It is unclear whether predators affect fitness of both sexes directly, by reducing survival, or indirectly, by affecting mate-searching. We empirically examined effects of increased predation risk on mate-searching behaviour and survival of male and female tree crickets, and their effects on mating success, using field-enclosure experiments with tree crickets *Oecanthus henryi* and their primary predator, green lynx spiders, *Peucetia viridans*. Crickets were allocated into three treatments with differential levels of predation risk. Increased predation risk strongly reduced survival, and thereby mating success, for both sexes. With increasing predation risk, males reduced calling and increased movement towards neighbouring callers, which had negative effects on mating success. Using simulations, we found male movement was significantly directed towards other calling males implying a switch to satellite strategies. Female movement behaviour, however, remained unaltered. Males and females thus differed in their response to comparable levels of predation risk, showing that the role of predation as a driver of alternative mate search strategies is sex and strategy-specific.

## Introduction

Predation plays an important role in shaping prey behaviour, with consequences for foraging, reproduction and other fitness-influencing activities. Individuals commonly acquire mates with reproductive strategies that vary across and within species (Gross 1996; Brockmann 2001; Oliveira et al. 2008). Prey that signal to attract mates risk being conspicuous to predators and parasitoids (Zuk and Kolluru 1998). Predation and parasitism have been hypothesised to be a potential cause for evolution of alternative reproductive tactics (ARTs), which are variable, often discontinuous, behaviours that allow a particular sex to achieve mating success and are observed in many animal species (Cade 1975; Gross 1996; Brockmann 2001; Oliveira et al. 2008). Several potential causes for the evolution and maintenance of ARTs have been hypothesised including intrasexual competition, mate choice (sexual selection) and predator evasion (natural selection) (Oliveira et al. 2008).

Some of these drivers of ARTs in mate searching and courting have been explored in crickets and frogs (Cade 1975; French and Cade 1989; Cade and Cade 1992; Lucas and Howard 1995; Walker and Cade 2003; Lucas and Howard 2008; Castellano et al. 2009; Rotenberry et al. 2015). Males of some species either call and attract females or silently search and intercept them, and these are known as calling and satellite strategies, respectively (Wells 1977; Cade 1979). Caller/satellite strategies in crickets are a classic system to theoretically and experimentally study the causes and consequences of ARTs (Cade 1975; French and Cade 1989; Cade and Cade 1992; Walker and Cade 2003). Relative fitness of caller/satellite strategies under varying conditions such as differential densities, sex ratios and body size have been tested either empirically (Cade 1975; French and Cade 1989; Cade and Cade 1992; Castellano et al. 2009) or in theoretical models (Rowell and Cade 1993; Lucas and Howard 1995; Walker and Cade 2003; Lucas and Howard 2008; Rotenberry et al. 2015). For instance, empirically varying cricket densities revealed more calling at lower densities, which was also correlated with higher mating success (French and Cade 1989; Cade and Cade 1992). Theory predicts relative fitness of ARTs in the calling-satellite system to be dependent on varying parasitism risk (Walker and Cade 2003; Rotenberry et al. 2015). Unlike these drivers of ARTs, however, predation as a potential cause of ARTs in the mate searching context has been less explored empirically.

Prey are known to mitigate predation risk by modifying their behaviours (reviewed in (Lima 1998; Preisser et al. 2005)). Considering sexually attractive signalling for instance, guppies minimise risk of predation on the visually conspicuous courtship behaviour by exhibiting sneak copulation tactics (Endler 1987). Similarly, signalling males of tree frogs and Tungara frogs reduce their calling effort in the presence of model predators (Tuttle and Ryan 1982; Ryan 1985). Increasing predation risk can thus affect fitness of prey species not only by reducing their survival, but also by scaring prey into modifying fitness-influencing behaviours. The few studies investigating predation risk as a driver of ARTs in the context of courtship (Wilgers et al. 2014) and territoriality (Oliveira et al. 2016) have not, however, examined the direct effect of predation in the form of mortality. We therefore investigated whether varying predation risk affects prey fitness due to reduced survival or altered reproductive tactics in the context of mate searching. With increased risk of predation, crickets are predicted to employ less conspicuous ARTs to search for and acquire mates (Walker and Cade 2003; Rotenberry et al. 2015).

ARTs have been predicted to evolve more frequently in males due to higher maximum potential benefits from multiple matings in males and higher investment in offspring by females (Taborsky et al. 2008). However, ARTs have also typically been studied in males (Brockmann 2008) and recent work on female competition has revealed female ARTs as well (Brockmann 2001; Johnson and Brockmann 2012; Hill et al. 2015; Ferrari et al. 2019). In prey systems where both males and females contribute to mate search, it is thus crucial to explore responses of both sexes to predation risk. For example in guppy courtship, increasingly successful sneaky copulation attempts by males in the presence of predation risk (Endler 1987) were later found to be better explained by predator-driven reduction in female preference (Magurran and Seghers 1994; Godin and Briggs 1996). No studies have, however examined ARTs simultaneously in both sexes of a species in the mate search context. In parallel with studying calling in males, we therefore also tested movement in females as a fitness-influencing behaviour that could change in response to varying predation risk.

We conducted field-enclosure experiments across several nights under variable predation risk, to address the following objectives: 1) With increasing predation risk, do male crickets shift from calling to satellite behaviour and do females alter their movement behaviour? 2) Does increasing predation risk reduce survival in male and female crickets? 3) Is mating success better predicted by survival, or by the extent and kind of mate-searching behaviours? We expected both male and female crickets to switch to less risky ARTs and their survival to decrease with increasing predation risk, with both changes consequently influencing fitness.

## Methods

We carried out this study on a tree cricket species, *Oecanthus henryi* and its primary predator, the green lynx spider, *Peucitia viridans* (Torsekar et al. 2019). *Oecanthus henryi* is observed on the bushes of *Hyptis suaveolens* in the dry scrublands of southern India. Males produce long-distance acoustic calls to which silent females respond by localising them, in the context of pair formation (Walker 1957; Bhattacharya 2016). We conducted field-enclosure experiments in a field of homogeneously distributed *H. suaveolens* near Ullodu village (13°38’27.2“N 77°42’01.1”E) in the Chikkaballapur district of Karnataka state in southern India, from February 2016 to May 2017. We set up these experiments inside enclosures of dimensions 6 m × 6 m × 2.2 m, constructed using wooden stakes and fastened with a stainless-steel mesh (mesh size: 0.1 cm × 0.2 cm). The enclosures were constructed around naturally growing *Hyptis suaveolens* bushes. We tagged and numbered all bushes inside the enclosures and ensured the densities and characteristics of bushes were comparable. We maintained three predation risk treatments that differed in the number of spiders released inside the enclosure. ‘No predation’ treatment (N = 2) involved no spiders or other predators present inside the enclosure, whereas ‘low predation’ (N = 3) and ‘high predation’ (N = 3) treatments included 15 spiders and 120 spiders, respectively. Fifteen male and fifteen female crickets were released in each enclosure for all treatments. We marked crickets with unique tricolour codes using non-toxic paint markers (Edding 780, Edding, St Albans, U.K.) to distinguish individuals. A trial consisted of monitoring male and female crickets for seven consecutive nights.

Spiders were sized to confirm that they were large enough to be predators of crickets (details in (Torsekar et al. 2019)). Prior to the commencement of every trial, all bushes inside the enclosures were carefully inspected and any adults or nymphs of *O. henryi*, *P. viridans* or any other potential predators of *O. henryi* found were caught and released outside the enclosures. 24 hours prior to the start of each trial, we released marked crickets inside the enclosures on randomly chosen bushes, to let them acclimate. Spiders were released on randomly chosen bushes an hour before the start of the trial. In accordance with the methodology of Walker and Cade (2003), we did not add new crickets through the duration of a trial to replace dead crickets. Accordingly, as the number of crickets reduced, we removed spiders every alternate night to maintain the ratio of crickets to spiders that determined the predation risk treatment. Once a trial began, we recorded the location and behaviour of crickets from 1900 hours to 2130 hours at two temporal resolutions. Location of individual crickets and whether there was a spider co-occurring with them on the same bush was recorded every 60 minutes, thrice a night. We recorded whether or not a male was calling or mating every 10 minutes, i.e. 16 times per night. Mating durations in *O. henryi* typically range from 20-45 minutes (Deb 2015). During each 10-minute scan, males were identified visually and acoustically and whether they were calling or mating was noted. We also noted the bush on which each cricket was found.

### Measure of predation risk

We measured predation risk encountered by crickets by quantifying spatial proximity between crickets and spiders. Co-occurrence of both on a bush was used as a non-arbitrary definition of spatial proximity. For an individual cricket, we collated all such co-occurrences across all sampling points during which it was observed, to calculate its probability of co-occurrence with a predator on a bush. This provided a finer resolution and more accurate measure of risk, at the individual level, rather than considering treatment as a categorical measure of risk faced by a population of crickets.

### Measure of mate searching behaviour: Females

We measured inter-bush movement of female crickets between sampling points within nights using two metrics: (a) the distance moved and (b) the likelihood of such movement. Successive sampling points were considered only within a night, discounting any change in location across nights. For measuring distance moved within nights, we used Euclidean distances between bushes. Euclidean distances were calculated before trials began, by carefully recording spatial location of every bush in each enclosure as polar co-ordinates, using a reference point common to each enclosure (Survey Compass 17475780, error ±0.5°, conceptualized by Francis Barker and Sons Ltd., sold and serviced by Lawrence and Mayo, India). This information along with the location data of each individual was used to measure the distance moved by each female cricket. Movement was also analysed as likelihood of movement to examine whether crickets move more often as a function of varying predation risk, regardless of how much they moved.

### Measure of mate searching behavior: Males

Calling in male crickets was measured in two ways, (a) how much a male called (calling effort) and (b) likelihood of calling. Calling effort was measured as the proportion of scans in which each male was observed calling. Calling effort of a male cricket was therefore the total sampling points it was found calling divided by the total sampling points it was scanned and not found mating. We measured likelihood of calling as whether a male called or not on a given night, regardless of how much it called. Movement of males was measured in the same way as for females (described in earlier section).

### Satellite behaviour

Increased movement may suggest a shift in male mate-searching from signaling to satellite behaviour. But alternative explanations for this behavioural modification are plausible. For instance, males may move more to escape spider attacks or to search for females without performing satellite behaviour. We explored what drives movement in males by investigating whether their movement implied satellite behaviour. Every time a male moved across bushes, we listed all calling males it could hear before it moved (hearing threshold based on (Deb et al. 2012)). Distances between the locations of these calling males and the focal male’s new location were noted, as well as the distance to the nearest calling neighbour. To understand if this is directed movement, the focal male’s new location was simulated (5000 times, to any of the available bushes in the enclosure), distances between each simulated location and all callers it could hear before it moved were noted, and all nearest neighbour distances were recorded. The median of this distribution was defined as the simulated nearest neighbor distance in the absence of orientation towards another calling male. We ran these simulations with a manually written ‘for loop’ code in R software version 3.3.3 (Team 2018) (for code details refer to supplementary information S2). We interpreted whether movement is directed towards a caller, implying satellite behaviour, by comparing the real and simulated nearest neighbour distances to a caller, using permutation tests.

### Measure of survival and mating success

Survival was measured as the number of nights each individual survived. Crickets were counted thrice every night. Since dead bodies of crickets were almost never found (ants immediately carried any small carcasses away), all crickets missing for more than one night and which did not reappear on subsequent nights were recorded as dead. We recorded mating success as the number of matings an individual acquired. One mating event was counted when crickets were found mating in at least one scan.

### Statistical analyses

For validating our treatments, we compared the co-occurrence probabilities of crickets and spiders on the same bushes across the duration of the experiment, in different treatments. We ran pair-wise permutation tests between the three treatments to test whether the probability of co-occurrences of crickets in each treatment were statistically different from each other based on *P*-values. These analyses were also run separately for male and female crickets.

We hypothesised that varying predation risk would affect mate searching behaviour and survival in crickets, which would consequently affect their mating success. We first tested change in survival and mate searching behaviour as a function of varying predation risk in separate models. Specifically, we examined the effect of varying predation risk on the following response variables; for females: survival, distance moved and likelihood of movement, and for males: survival, calling effort, likelihood of calling, distance moved and likelihood of movement. Next, we explored the effect of survival and mate searching behaviour on mating success, separately for male and female crickets.

We analysed the effect of predation risk on mate searching, separately for male and female crickets. For this analysis, we considered each night for every individual as a separate data point (Males: N = 506; Females: N = 479). Such fine resolution allowed for a better understanding of how individuals behave depending on the risk they face per night. We used zero-inflated negative binomial GLMMs for analysing distance moved by male and female crickets and binomial GLMMs for calling effort, likelihood of calling and likelihood of movement. We ran an additional analysis for examining how likelihood of movement interacts between the sexes with increasing nightly predation risk. Co-occurrence probabilities with spiders was the fixed effect and individual ID was the random effect for all models testing mate searching behaviour. We tested how survival changed due to predation risk for each individual cricket over the duration of the experiment using Poisson GLM with co-occurrence probabilities and sex and their interaction as the predictors (Males: N = 113; Females: N = 111). Mating success of individuals was analysed as a function of how long individuals survived and how they communicated, separately for the sexes, using Poisson GLMs (Males: N = 113; Females: N = 110). For all analyses, non-significant interaction terms (*P* > 0.05) were dropped from the model. We calculated *P*-values by running permutation tests for statistical hypothesis testing (Manly 2018) and also calculated effect sizes and their associated 95% confidence intervals (CI) (Nakagawa and Cuthill 2007). For interpretation, 95% CI of regression coefficients not overlapping with zero was considered to have ‘strong support’ for predictions; 95% CI slightly overlapping zero, up to 85% CI were considered to have ‘moderate support’, and greatly overlapping with zero were regarded to have ‘no support’ (Cumming 2013; Abbey-Lee et al. 2016). All analyses were run in R software version 3.3.3 (Team 2018). Data collation and manipulation were done using the dplyr package (Wickham et al. 2017) and visualisation using the ggplot package (Wickham 2009) (for details refer to supplementary information S1).

## Results

### Predation risk

Crickets in each treatment faced significantly different co-occurrence probabilities with spiders (*P* < 0.001; Fig. 1a). These results were corroborated when co-occurrence probabilities of male and female crickets with spiders in different treatments were tested separately (*P* < 0.001). The two sexes did not face different co-occurrence probabilities within each treatment (*P* = 0.99; Fig. 1b). Therefore, crickets experienced different predation risk across treatments, whereas the two sexes experienced similar risk within treatments.

**Figure 1.**
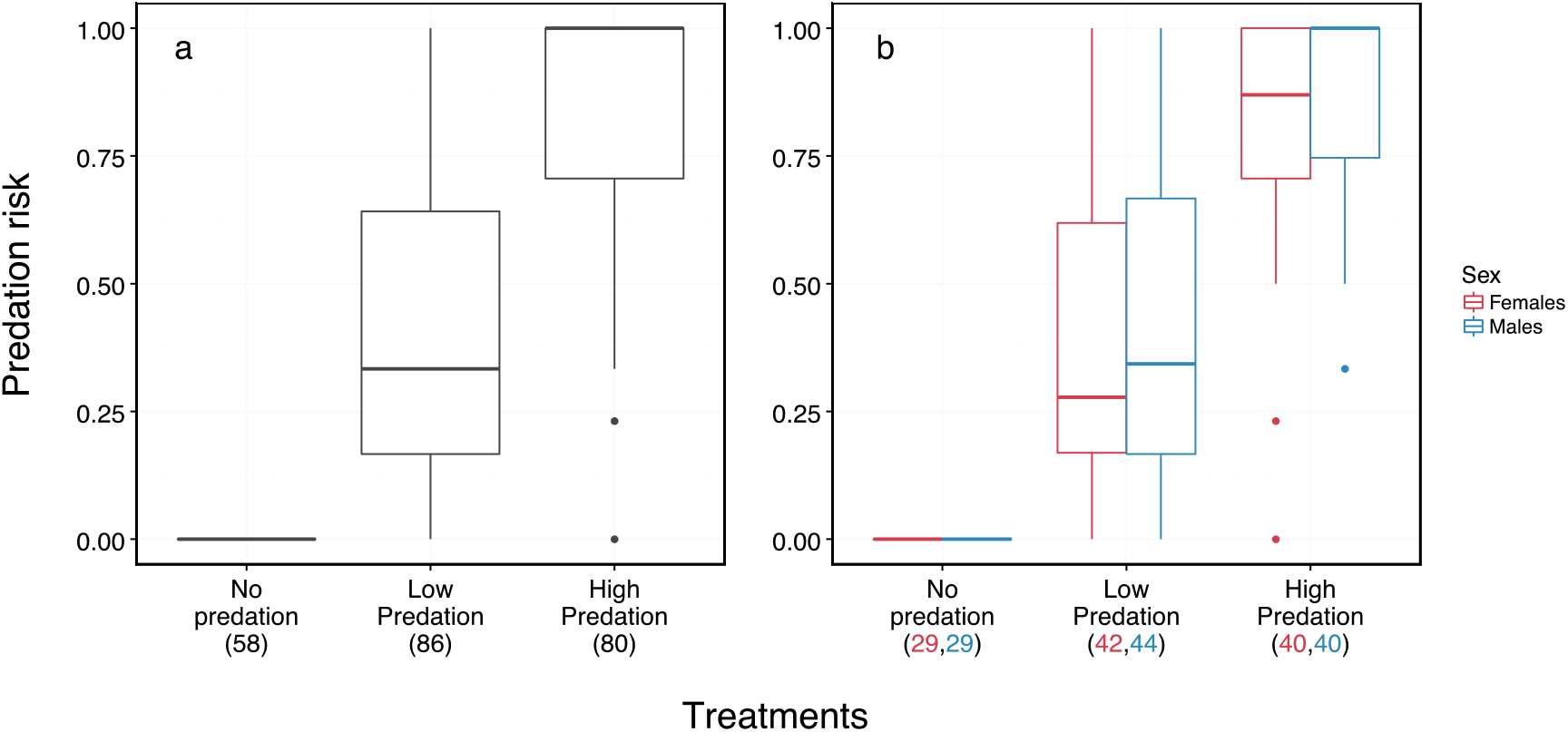
Predation risk encountered by crickets in different treatments. This comparison is shown for a) all crickets in the respective treatments and b) separately for male and female crickets. Predation risk is the probability of co-occurrence of a cricket with a spider on a bush across all nights it survived till the end of the experiment. Numbers in parentheses are sample sizes (number of crickets).

### Mate searching behavior

Predation risk affected mate searching behaviours in male crickets (Table S1, supplementary material). There was strong support for males calling less (χ^2^ = 73.790, *P* < 0.001; Fig. 2a) and weak support for decreased likelihood of calling (χ^2^ = 3.181, *P* = 0.074; Fig. 2b). Increasing nightly predation risk increased the likelihood of males moving across bushes (χ^2^ = 4.772, *P* = 0.029; Fig. 2d) but did not significantly affect how far males moved (χ^2^ = 0.291, *P* = 0.59; Fig. 2c). Calling effort of males was on average 15% less when there was a spider present on the same bush in comparison with when there was not. In female crickets, neither distance moved nor likelihood of movement showed association with increasing predation risk (Fig. 2e and 2f; Table S2). With increasing predation risk, there was no evidence for change in within-night distance moved (χ^2^ = 0.174, *P* = 0.677; Fig. 2e) or likelihood of movement (χ^2^ = 0.426, *P* = 0.514; Fig. 2f). Therefore, with increasing predation risk, males reduced how much they called and increased likelihood of movement, but movement by females was unaffected. Examining movement of both sexes together, female crickets showed higher likelihood of movement than males when spiders did not co-occur on the bush (Table S3).

**Figure 2.**
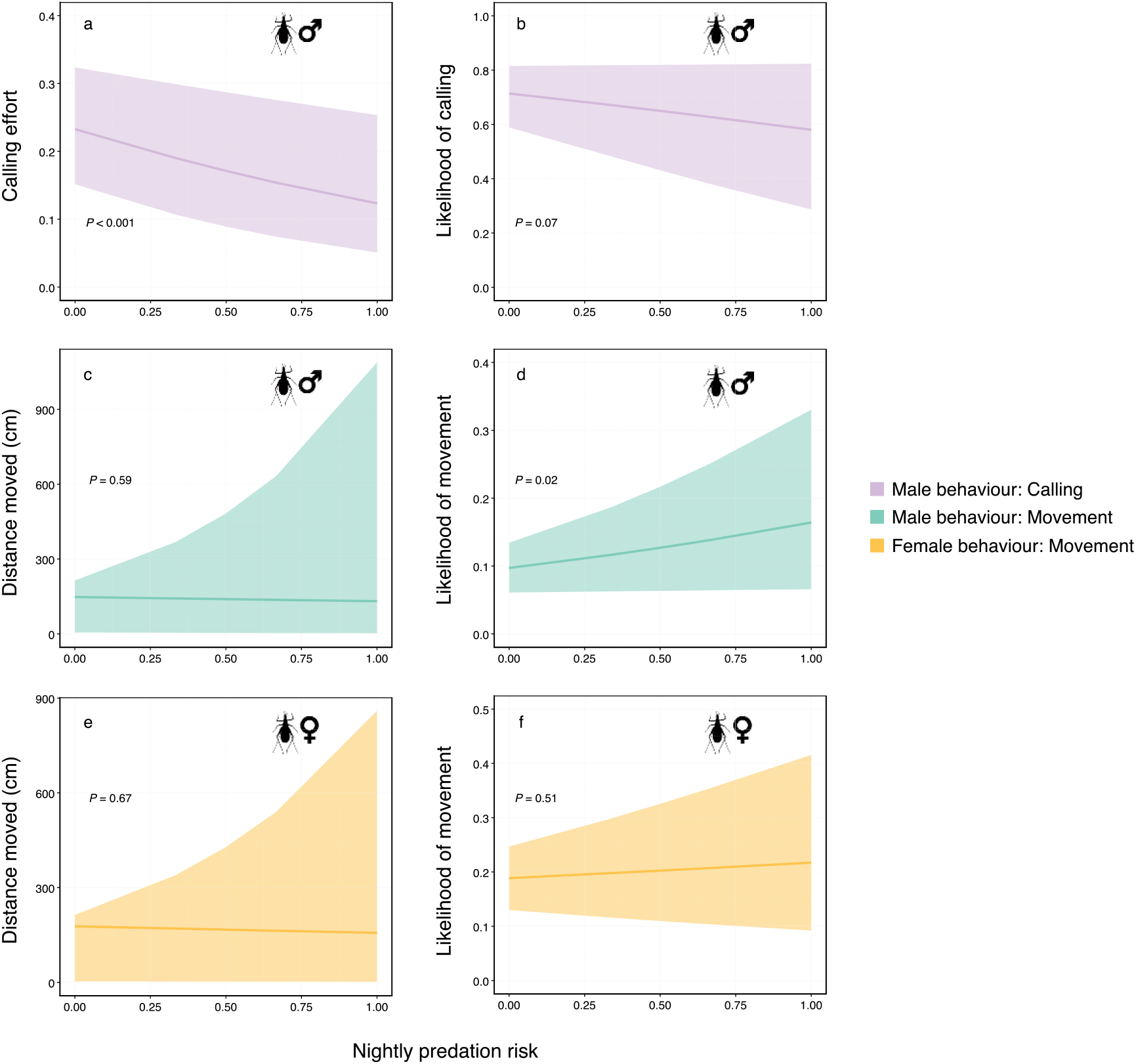
The effect of increasing predation risk on male and female behaviour including a) the calling effort per night; b) likelihood of calling; c) distance moved within a night (males); d) likelihood of movement (males); e) distance moved within a night (females); f) likelihood of movement (females). All X axes show predation risk faced by crickets, represented as the probability of co-occurrence with spiders per night. The lines are predictions based on GLMMs, with shaded areas representing bootstrapped 95% CI.

Nearest neighbour distances between new locations of moving males and calling males they could hear before they moved were significantly lower for the empirical data when compared with data simulating random movement of males (N = 62, *P* < 0.001; Fig. 3). Male cricket movement was therefore significantly directed towards calling males that they could hear around them.

**Figure 3.**
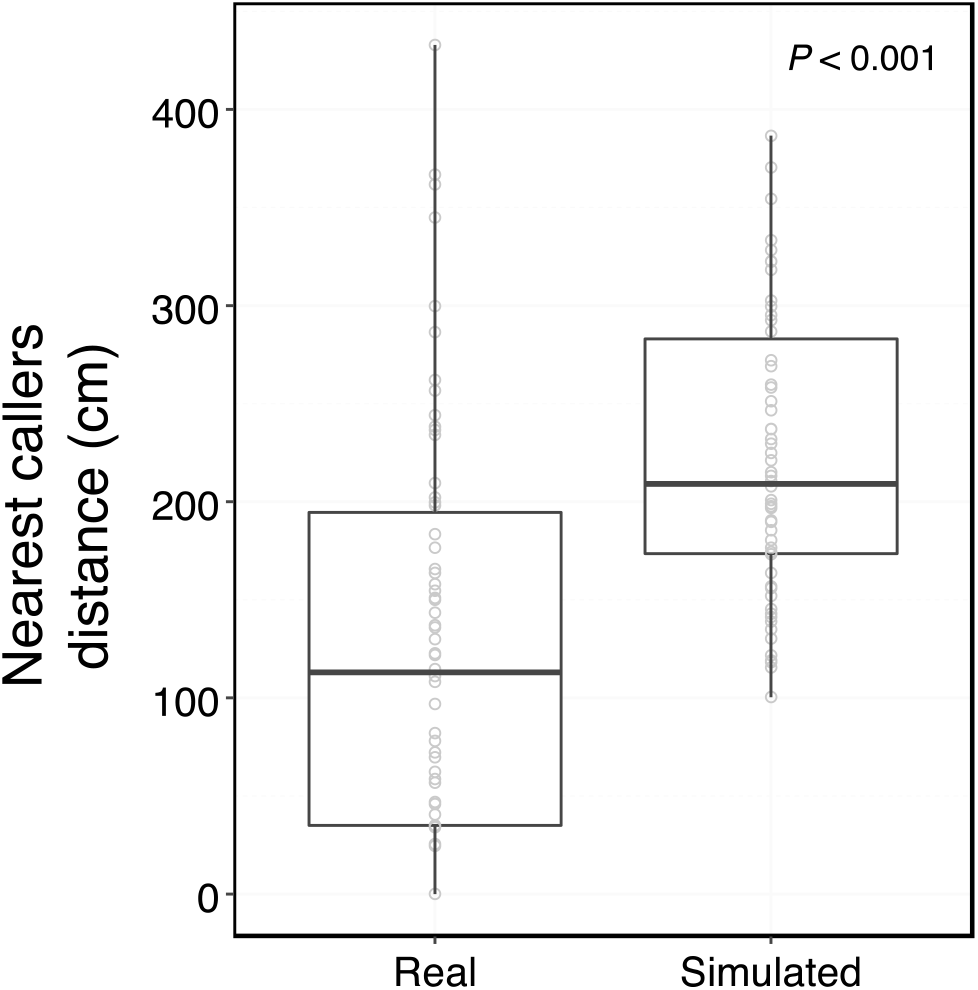
Movement as a satellite strategy. A comparison of distance to the closest caller when a male cricket moved in the enclosure. ‘Real’ represents the empirical data of male movement and ‘Simulated’ represents the simulated data based on the null hypothesis of random movement to any bush. Points shown are actual distances to closest caller in the ‘real’ category, and medians of distributions of distances to closest caller when movement was simulated.

### Survival

Survival of both male and female crickets significantly reduced with increasing predation risk (Table S4). This strong decrease in survival (χ^2^ =22.865, *P* < 0.001; Fig. 4; Table S4) was similar for both sexes (χ^2^ =1.796, *P* = 0.407; Fig. 4; Table S4).

**Figure 4.**
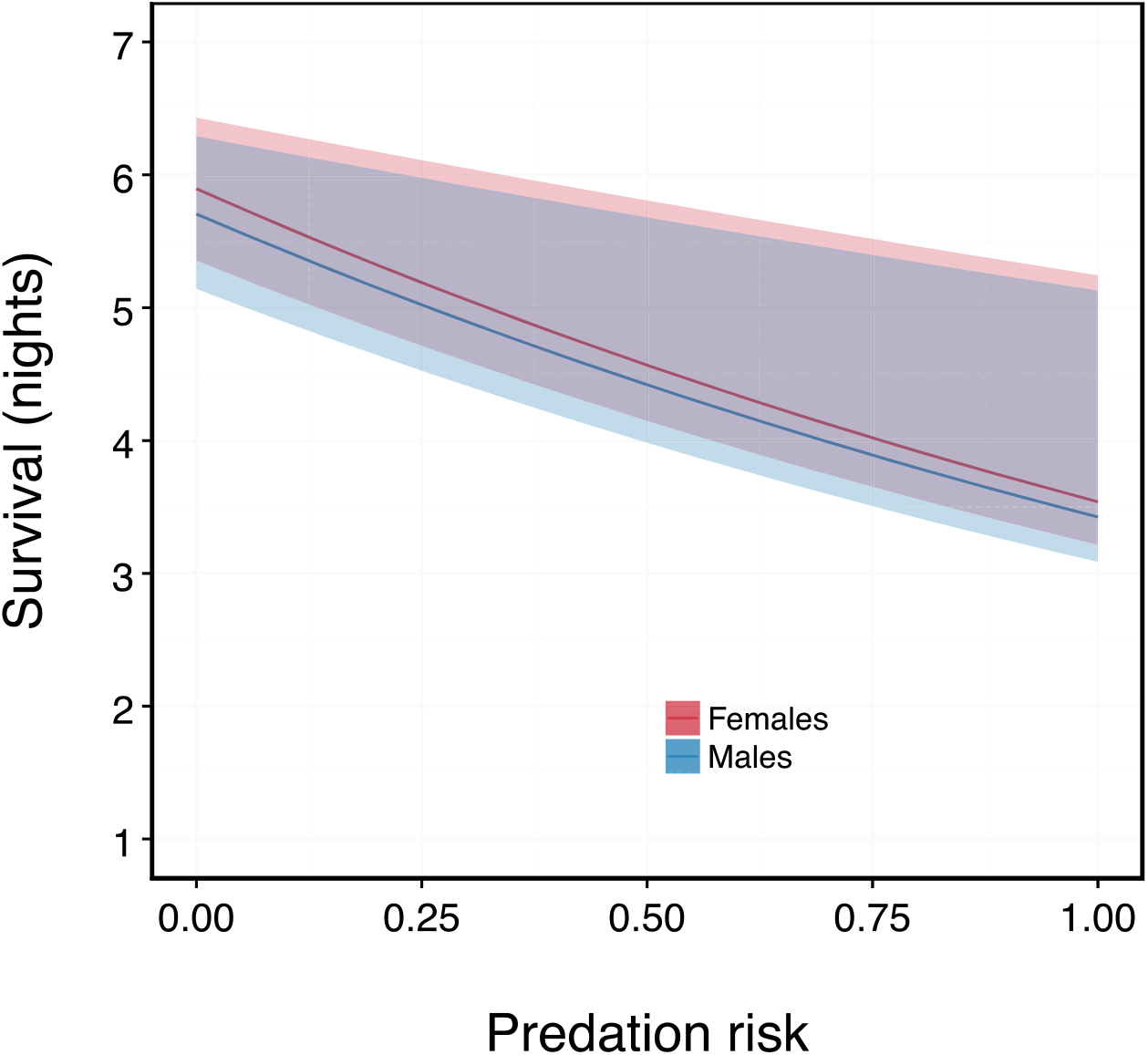
Survival as a function of predation risk for male and female crickets. The X axis is the predation risk faced by crickets, represented by the probability of co-occurrence with spiders on bushes across the nights they survived. The lines are predictions based on GLMs, with shaded areas representing bootstrapped 95% CI.

### Mating success

Crickets varied widely in the number of matings they acquired during the duration of the experiment (Males: 0.57 ± 0.83; Females: 0.57 ± 0.90, mean ± SD). To determine what best predicted this variation in total number of matings, we tested individual mating success as a response to propensity to mate search, and survival, separately for male and female crickets. Mating success in male crickets (Table 1) increased with increasing survival (χ^2^ = 18.982, *P* < 0.001), decreased with increasing likelihood of movement (χ^2^ = 4.758, *P* = 0.029) and showed no association with calling effort (χ^2^ = 1.310, *P* = 0.252) when analysed together. Female mating success (Table 1) increased with increasing survival (χ^2^ = 12.111, *P* < 0.001) and was not influenced by how far individuals moved (χ^2^ = 0.049, *P* = 0.824).

**Table 1.** Mating success in male and female crickets, tested separately using Poisson GLMs. Model coefficients, bootstrapped 95% CI for coefficients and *P* values from permutation tests (based on 10000 iterations) are shown.

| *Term* | *Coefficient* | *95% CI* | $\chi^2$ | *P* |
| --- | --- | --- | --- | --- |
| **Males** |  |  |  |  |
| *Intercept* | -2.306 | -3.600 – -1.547 |  |  |
| *Survival* | 0.312 | 0.169 – 0.518 | 18.982 | < 0.001 |
| *Calling effort* | 0.716 | -0.464 – 1.871 | 1.310 | 0.252 |
| *Likelihood of movement* | -0.381 | -0.797 – -0.068 | 4.758 | 0.029 |
| **Females** |  |  |  |  |
| *Intercept* | -1.891 | -3.443 – -1.004 |  |  |
| *Survival* | 0.238 | 0.076 – 0.470 | 12.111 | < 0.001 |
| *Distance moved* | 0.00011 | -0.0012 – 0.0008 | 0.049 | 0.824 |

## Discussion

With increasing predation risk, male crickets altered their mate searching tactic from calling to satellite behaviour, which had negative consequences on mating success. Females, however, did not exhibit ARTs, as predicted by theory (Taborsky et al. 2008). Survival of both was strongly affected by increasing predation risk but did not differ between the sexes. Although previous studies have simulated varying parasitism risk and predicted variable fitness for calling and satellite behaviours (Walker and Cade 2003; Rotenberry et al. 2015), to the best of our knowledge, this is the first empirical demonstration of these predictions.

Measuring the probability of crickets and spiders co-occurring on a bush as representative of predation risk facilitated the validation of predation treatments in which different samples of crickets experienced disparate risk of predation. Previous work in this system has shown probability of co-occurrence to be a reliable indicator of predation risk (Torsekar et al. 2019). Our setup enabled us to measure predation risk faced by individual crickets empirically, allowing examination of prey responses as a function of the risk encountered by individuals per night, and resultant individual fitness consequences.

Increasing predation risk not only reduced calling effort and likelihood of calling in male crickets, but also simultaneously increased their likelihood of movement, indicating that males might be employing the less conspicuous tactic of silent searching instead of calling to obtain mates. Reduced calling and increased movement directed towards neighbouring callers suggests a shift in male mate searching from signaling to satellite behaviour. Simulations of the caller-satellite system show that ARTs may be both frequency- and environment-dependent (Walker and Cade 2003; Rotenberry et al. 2015). Frequency-dependence in this system has been demonstrated empirically in field crickets (French and Cade 1989; Cade and Cade 1992). Our findings provide empirical evidence for environment-dependence, where males favoured satellite over calling behaviour under the condition of high predation risk. Other examples of ARTs being affected by frequency, along with environment-dependence include bluegill sunfish (Gross 1997) and horseshoe crabs (Brockmann and Penn 1992).

Female mate searching behaviour was not affected by varying predation risk; neither the likelihood of moving nor how much they moved. A potential explanation is that female propensity to move is sufficient to escape predation. Female likelihood of movement, regardless of predator presence, is comparable with male likelihood of movement when a spider is present on the bush (Table S3), suggesting that female movement is enough to avoid capture. But do female crickets exhibit ARTs while mate searching? A study found that only 30% of wild-caught female crickets exhibited phonotaxis, suggesting variation in mate searching behaviour (Torsekar et al. 2019). Importantly, females not performing phonotaxis are motivated to mate when presented an opportunity (Modak, Brown and Balakrishnan, unpublished results), implying female ARTs in mate searching behaviour (Johnson and Brockmann 2012). Therefore, although not dependent on environmental perturbations such as predation risk, female ARTs may be frequency- or condition-dependent. For example, mating status is a strong driver of phonotaxis behaviour, and hence responsiveness, in *O. henryi* females (Modak, Brown and Balakrishnan, unpublished results). This suggests that ARTs in female mate searching behaviour may be influenced by intrasexual and intersexual rather than natural selection mechanisms. Overall, male and female crickets responded differently to varying predation risk, despite experiencing similar predation risk.

Survival and satellite behaviour influenced mating success in males, whereas survival alone explained mating success in female crickets. Survival was expected to have a strong effect on mating success and this is consistent with the simulation study results, in which the number of encountered females was strongly correlated with the lifespan of male crickets (Walker and Cade 2003; Rotenberry et al. 2015). Contrary to our expectations, however, calling effort did not on its own influence mating success. A possible explanation is that since survival and calling effort show a weak positive correlation (r = 0.33), callers are represented by individuals that survive for longer. Studies have shown that expression of sexually selected traits in males is positively correlated with their age (reviewed in (Jennions et al. 2001); but see (Hunt et al. 2004)). In other words, males that survive for long and increase their chances for achieving mating success, also exhibit higher calling effort. On the other hand, we find that with increasing likelihood of movement behaviour, mating success reduced. We speculate that male crickets switch from calling to satellite behaviour when predation risk is high because the satellite/search strategy is safer. Even though mating opportunities are affected, the switch to satellite strategy may allow males to survive for long enough to mate later.

In conclusion, our results provide the first empirical demonstration that males change their mate search behaviour under increased predation risk conditions whereas females, under equivalent conditions of risk, do not. Also, fitness consequences of response to immediate environmental threats are different for the two sexes, with only direct (survival) effects in females, but both direct and indirect (mate searching behaviour) effects in males. Taken together, these are likely to have important implications for population-level selection processes, further emphasizing the importance of studying ARTs in both sexes.

## Acknowledgements

We thank Kada Reddy and Manjunatha Reddy for their help in constructing the enclosure and Prakash, Chandra, Nanjundaiah and Manjunatha for their assistance in fieldwork. We appreciate all the helpful discussions with Diptarup Nandi, Kavita Isvaran, Manvi Sharma and Ian Durbach during the research. We thank Vivek Nityananda and Aparna Lajmi for their comments on the manuscript.

## Funding

We thank the DBT-IISc Partnership Programme (phase 1 and 2) of the Department of BioTechnology (DBT), Government of India for funding the fieldwork, DST-FIST (Dept. of Science & Technology Fund for Improvement of S & T Infrastructure, Govt. of India) for some of the equipment used in the study and the Ministry of Human Resource Development, Government of India for the research fellowship of VRT.

## Author contributions

VRT participated in conceptualising and designing the study, carried out data collection and analysis and wrote the manuscript. RB contributed to conceptualising and designing the study, interpretation of data and writing the manuscript.

## Ethical approval

All behavioural data sampling and experiments were performed in accordance with the national guidelines for the ethical treatment of animals laid out by the National Biodiversity Authority (Government of India)

## Notes

https://github.com/torsay/predation_risk_mate_searching/blob/master/satellite_simulations

