## Supplementary information for "Sex differences in alternative reproductive tactics in response to predation risk in tree crickets"

**S1 Details of statistical analysis**

*Mate searching behaviour*

We analysed the effect of predation risk on mate searching, separately for male and female crickets. Distance moved by male and female crickets were non-normal continuous data that were zero-inflated and overdispersed. Thus, data were analysed using the zero-inflated negative binomial GLMM in the glmmTMB package (Brooks et al. 2017) in R. Since calling effort, likelihood of calling and likelihood of movement are proportions bounded between 0 and 1, binomial GLMMs were used to analyse the data using the lme4 package (Bates et al. 2014). Repeated observations of individual crickets were accounted for by including individual ID as a random effect. For all analyses, non-significant interaction terms (*P* > 0.05) were removed from the model.

*Survival*

We tested how survival changed with varying predation risk for each individual cricket over the duration of the experiment. To compare and interpret results of male and female crickets, survival of both were analysed in the same model. Whether survival was affected by varying predation risk depending on the sex of the cricket was tested by including a two-way interaction term. We ran a GLM assuming Poisson-distributed errors since survival data were non-normal counts with comparable mean and variance. Predation risk, the single predictor, was represented by co-occurrence probabilities of individuals with spiders which were collated across the number of nights they survived.

*Mating success*

Mating success of individuals was analysed as a function of how long individuals survived and how they communicated. For male crickets, only calling effort and likelihood of movement were considered in the model, since including likelihood of calling and distance moved were collinear with the chosen variables (Zuur et al. 2009). For similar reasons, only distance moved was considered as an explanatory variable for females. We used variance inflation factors (VIF) to assess which explanatory variables are collinear and should be dropped (Zuur et al. 2009). VIF values of our final models were below 1.5, implying that multi-collinearity is not a concern (Zuur et al. 2009). GLM assuming Poisson-distributed errors were run because mating success data were non-normal counts. Since densities of crickets were not maintained through the duration of the experiment, potential effects of resultant differential encounter probabilities between the sexes were tested in the model and dropped when found to be not significant.

We calculated *P*-values by running permutation tests for statistical hypothesis testing (Manly 2018) and also calculated effect sizes and their associated 95% confidence intervals (CI) (Nakagawa and Cuthill 2007). This provided evidence for uncertainty around regression coefficients and helped bolster our inferences. We measured non-parametric 95% CI by bootstrapping model coefficients, by re-sampling with replacement 10,000 times, while maintaining the random effect structure for mixed models (Manly 2018).

**S2 Code for validating satellite behaviour using simulations**

We explored what drives movement in males by investigating whether their movement implied satellite behaviour. We employed simulations to better understand whether male movement was directed towards calling males. The code written in R software is made available at https://github.com/torsay/predation_risk_mate_searching/blob/master/satellite_simulations

**Table S1**: Generalised linear mixed-effects models fitted to analyse male mate searching behavior over increasing nightly predation risk (N = 506). Calling effort, likelihood of calling and likelihood of movement were analysed using binomial GLMMs, and distance moved by males per night was analysed using zero-inflated poisson GLMM. Model coefficients, bootstrapped 95% CI for coefficients and *P* values from permutation/randomization tests (based on 10000 iterations) are shown.

|  | **Calling effort** | | | | **Likelihood of calling** | | | |
| --- | --- | --- | --- | --- | --- | --- | --- | --- |
|  | **Coefficient** | **95% CI** | **χ^2^** | ***P*** | **Coefficient** | **95% CI** | **χ^2^** | ***P*** |
| Intercept | -1.194 | -1.723 – -0.738 |  |  | 0.913 | 0.359 –1.484 |  |  |
| Predation risk | -0.768 | -1.203 – -0.343 | 73.790 | < 0.001 | -0.590 | -1.270 – 0.055 | 3.181 | 0.074 |

|  | **Distance moved** | | | | **Likelihood of movement** | | | |
| --- | --- | --- | --- | --- | --- | --- | --- | --- |
|  | **Coefficient** | **95% CI** | **χ^2^** | ***P*** | **Coefficient** | **95% CI** | **χ^2^** | ***P*** |
| Intercept | 4.413 | 1.778 – 5.366 |  |  | -2.229 | -2.734 – -1.863 |  |  |
| Predation risk | -0.1408 | -0.640 – 1.626 | 0.291 | 0.590 | 0.601 | 0.0831 – 1.156 | 4.772 | 0.029 |

**Table S2**: Generalised linear mixed-effects models fitted to analyse female mate searching behavior over increasing nightly predation risk (N = 479). Likelihood of movement was analysed using binomial GLMMs and distance moved by females per night was analysed using zero-inflated poisson GLMM. Model coefficients, bootstrapped 95% CI for coefficients and *P* values from permutation/randomization tests (based on 10000 iterations) are shown.

|  | **Distance moved** | | | | **Likelihood of movement** | | | |
| --- | --- | --- | --- | --- | --- | --- | --- | --- |
|  | **Coefficient** | **95% CI** | **χ^2^** | ***P*** | **Coefficient** | **95% CI** | **χ^2^** | ***P*** |
| Intercept | 5.180 | 1.120 – 5.349 |  |  | -1.459 | -1.901 – -1.117 |  |  |
| Predation risk | -0.123 | -0.425 – 1.394 | 0.174 | 0.677 | 0.177 | -0.386 – 0.775 | 0.426 | 0.514 |

**Table S3**: Likelihood of movement in male and female crickets analysed together as a function of increasing predation risk using poisson GLMM. Model coefficients, bootstrapped 95% CI for coefficients and *P* values from permutation/randomization tests (based on 5000 iterations) are shown.

|  | **Coefficient** | **95% CI** | **χ^2^** | ***P*** |
| --- | --- | --- | --- | --- |
| Intercept (Females) | -1.488 | -1.830 - -1.183 |  |  |
| Predation risk | 0.358 | -0.023 - 0.743 | 13.074 | 0.592 |
| Sex: Males | -0.727 | -1.133 - -0.352 | 3.372 | <0.001 |

**Table S4**: Survival in male and female crickets analysed as a function of increasing predation risk using poisson generalised linear model. Model coefficients, bootstrapped 95% CI for coefficients and *P* values from permutation/randomization tests (based on 10000 iterations) are shown.

|  | **Coefficient** | **95% CI** | **χ^2^** | ***P*** |
| --- | --- | --- | --- | --- |
| Intercept (Females) | 1.774 | 1.679 – 1.860 |  |  |
| Predation risk | -0.356 | -0.509 – -0.204 | 22.368 | <0.001 |
| Sex: Males | -0.032 | -0.147 – 0.085 | 0.299 | 0.584 |

Brooks, M. E., K. Kristensen, K. J. van Benthem, A. Magnusson, C. W. Berg, A. Nielsen, H. J.

Skaug, et al. 2017. glmmTMB balances speed and flexibility among packages for zero-inflated generalized linear mixed modeling. The R Journal 9:378–400.

Manly, B. F. 2018. Randomization, bootstrap and Monte Carlo methods in biology. Chapman and Hall/CRC.

Nakagawa, S., and I. C. Cuthill. 2007. Effect size, confidence interval and statistical significance: a practical guide for biologists. Biological reviews 82:591–605.

Zuur, A. F., E. N. Ieno, N. Walker, A. A. Saveliev, and G. M. Smith. 2009. Mixed effects models and extensions in ecology with R. Springer Science & Business Media, New York, NY.
